# Infanticide by females is a leading source of juvenile mortality in a large social carnivore

**DOI:** 10.1101/2020.05.02.074237

**Authors:** Ally K Brown, Malit O Pioon, Kay E Holekamp, Eli D Strauss

## Abstract

Social animals benefit from their groupmates, so why do they sometimes kill each other’s offspring? Using 30 years of data from multiple groups of wild spotted hyenas, we address three critical aims for understanding infanticide in any species: (1) quantify the contribution of infanticide to overall mortality (2) describe the circumstances under which infanticide occurs and (3) evaluate hypotheses about the evolution of infanticide. We find that, although observed only rarely, infanticide is in fact a leading source of juvenile mortality. Infanticide accounted for 24% of juvenile mortality, and 1 in 10 hyenas born in our population perished due to infanticide. In all observed cases of infanticide, killers were adult females, but victims could be of both sexes. Of four hypotheses regarding the evolution of infanticide, we found the most support for the hypothesis that infanticide in spotted hyenas reflects competition over social status among matrilines.

## Introduction

Why do animals kill the offspring of their group members? Infanticide is a taxonomically widespread behavior, found in mammals, birds, reptiles, fish, and invertebrates (Agrell et al., 1998; Hausfater & Hrdy, 1984; Hrdy, 1979; O’Connor & Shine, 2004). In species where infanticide represents a common source of infant mortality, infant defense and avoidance of infanticidal individuals may function importantly in the developmental biology and social behavior of both adults and juveniles (Balme & Hunter, 2013; Lowe et al., 2018; Muller & Wrangham, 2002; Packer & Pusey, 1983). Thus, infanticide can play an important role in the biology of many species, provided that it occurs frequently enough to be an agent of selection. However, because mortality events occur infrequently and take place rapidly, the significance of infanticide is difficult to evaluate and remains unknown in many species. For instance, the existence of infanticide by male lions, now a canonical example of infanticide, was hotly debated as recently as the late 1990s (Dagg, 1998; Packer, 2000; Silk & Stanford, 1999).

Comparative studies have emphasized that a lack of observational data is a major barrier to the understanding of infanticide in ecology and evolution (Lukas & Huchard 2019). For species in which infanticide has not been specifically studied in detail, there is thus a critical need to (1) quantify the contribution of infanticide to overall mortality, (2) describe the contexts under which infanticide occurs, and (3) evaluate hypotheses about the evolution of infanticide. In this study, we address these needs in spotted hyenas (*Crocuta crocuta*), a large African group-living carnivore.

### Quantifying the contribution of infanticide to overall mortality

The first step to understanding the evolution of infanticide is to identify its contribution to mortality. In some species, infanticide contributes dramatically to overall mortality. For example, in one study, nearly a third of all African leopard offspring were killed by infanticide, suggesting strong selection pressure on juveniles and mothers to avoid infanticide (Balme & Hunter, 2013). No studies have yet investigated the frequency of infanticide in spotted hyenas, or how mortality due to infanticide compares to mortality from other sources. However, in a prior study, 48% of spotted hyenas perished in their first year of life, suggesting that juvenile mortality sources are strong agents of selection in this species (Watts & Holekamp, 2009). Furthermore, hyenas invest heavily in each offspring, bearing only litters of 1-2 juveniles, nursing them for over a year, and achieving an average lifetime reproductive success of only ~2 juveniles successfully reared to adulthood (Hofer & East, 2003; Strauss & Holekamp, 2019b). Taken together, this evidence suggests that infanticide might be an important behavior in this species if it represents a significant source of juvenile mortality.

### Describing the contexts in which infanticide occurs

Observational data describing infanticide are necessary to understand the impact of infanticide and its evolution (Lukas & Huchard, 2019), but such data for spotted hyenas are rare. Kruuk (1972) reported a case of attempted infanticide against a six-month old hyena and suggested that protection against infanticide by males was a potential driver of female dominance over males. In support of this view, a study of hyenas in the Serengeti documented two cases of males attempting infanticide against juveniles (East et al., 2003). Mills (1990) provided an alternative perspective on infanticide in spotted hyenas, suggesting that infanticide may be a form of inter-group competition, noting cases of cubs sustaining injuries or disappearing after hyenas from a neighboring clan visited the communal den. In a study of the uniquely masculinized genitals of female hyenas, Muller & Wrangham (2002) suggest that this unusual morphology may have evolved to protect female cubs against infanticide. In our study population, White (2005) reported a case of one female hyena killing two cubs (also reported here), and suggested that cubs may be at greatest risk of suffering infanticide right after being moved from the natal den to the communal den. This lack of consensus on infanticide in hyenas paired with the paucity of observational data reflects the need for more observational study of infanticide in this species.

### Evaluating hypotheses about the evolution of infanticide

Hrdy (1979) introduced the first adaptive hypotheses regarding the evolution of infanticide, positing that infanticide functions to provide fitness-related benefits to infanticidal individuals (Agrell et al., 1998). Hypothesized motivations to commit infanticide include the killing of conspecific offspring to increase the killer’s reproductive opportunities (‘sexual selection’ hypothesis), as a source of food (‘exploitation’ hypothesis), or to reduce intra-group competition for resources (‘resource competition’ hypothesis) (Hrdy, 1979).

Lukas & Huchard (2019) divide the resource competition hypothesis into four types, which vary in the societies in which they are expected to occur and the types of resources over which individuals compete. The ‘breeding space’ hypothesis suggests infanticidal individuals eliminate infants occupying spaces critical to reproduction, such as nest sites or dens. The ‘milk competition’ hypothesis suggests that infanticidal mothers kill unrelated offspring that attempt to nurse from them. The ‘allocare’ hypothesis posits that in cooperatively breeding species, dominant individuals maximize the number of helpers caring for their offspring by killing the offspring of subordinates within their group. Finally, the ‘social status’ hypothesis suggests that in stable societies, more dominant individuals kill the offspring of subordinate individuals to eliminate future rivals and protect the dominance status of themselves and their kin (Clutton-Brock & Huchard, 2013; Lukas & Huchard, 2019; Vullioud et al., 2019).

These hypotheses about the evolution of infanticide make different predictions about the circumstances under which infanticide is likely to occur. In our study of the evolution of infanticide in spotted hyenas, we ignore the ‘allocare’ and ‘milk competition’ hypotheses because hyenas aren’t cooperative breeders and because they have no trouble preventing unrelated offspring from nursing (Kruuk, 1972). We focus instead on the sexual selection, exploitation, social status, and breeding space hypotheses and their predictions (Table 1). The sexual selection hypothesis predicts that infanticide will be perpetrated by males and that mating opportunities should increase for males after they have committed infanticide. The exploitation hypothesis predicts that infanticide should occur when prey are scarce, and infanticidal individuals should consume the victim. Although hyenas do not compete with individuals from other groups over breeding space, they do share limited space in communal dens with their clan-mates. Thus, the ‘breeding space’ hypothesis predicts that infanticide should occur more frequently when there are many juveniles residing at the communal den. The ‘social status’ hypothesis predicts that infanticide should be perpetrated by high-ranking adults against the offspring of low-ranking individuals, and that the sex of the victims and perpetrators of infanticide should be biased towards the philopatric sex. Finally, it is worth noting that these hypotheses are not mutually exclusive, and it is possible that infanticidal individuals accrue multiple benefits from committing infanticide (Hrdy, 1979; Lukas & Huchard, 2019).

**Table 1.**
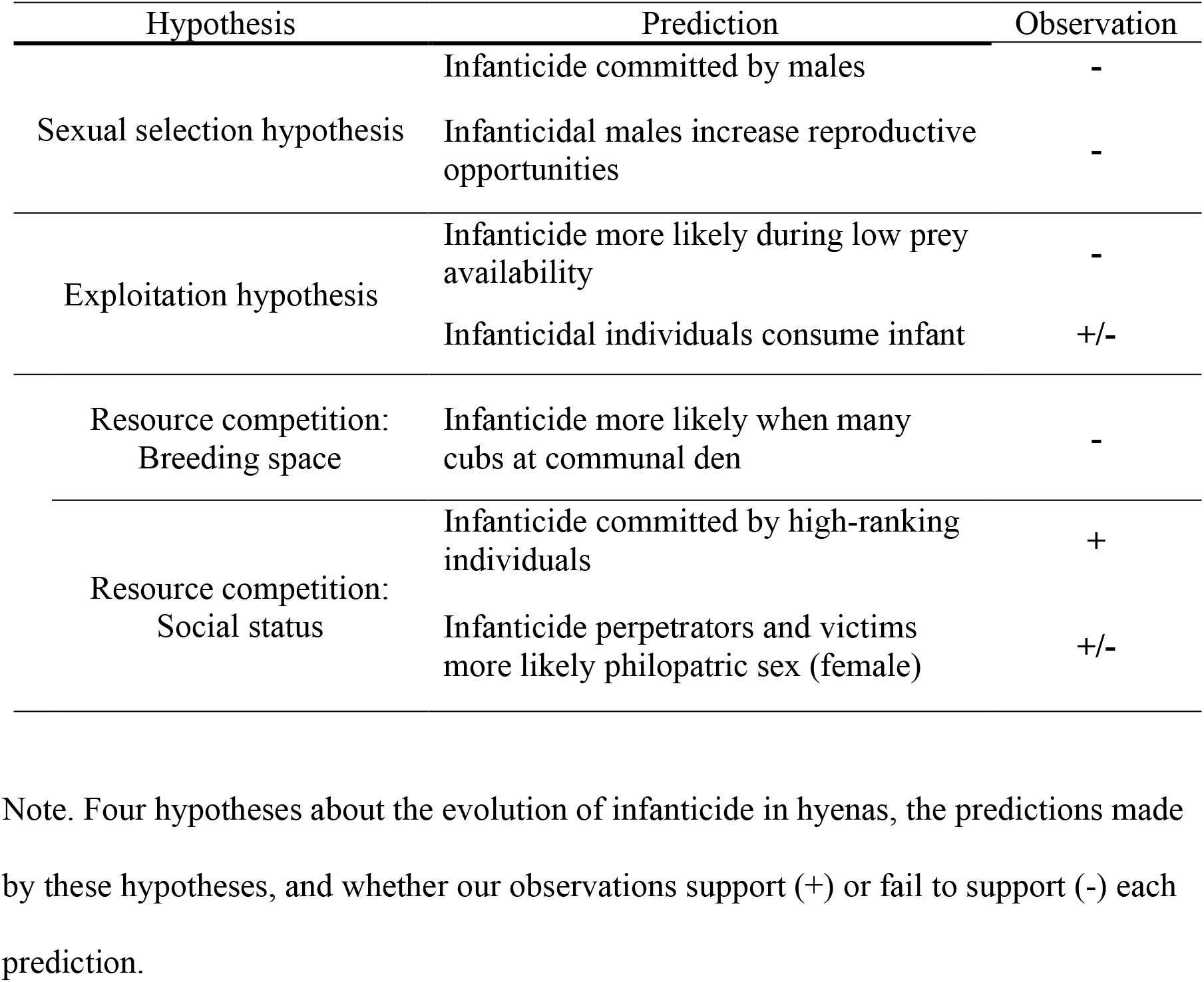
Hyena Infanticide Hypotheses

| Hypothesis | Prediction | Observation |
| --- | --- | --- |
| Sexual selection hypothesis | Infanticide committed by males | - |
|  | Infanticidal males increase reproductive opportunities | - |
| Exploitation hypothesis | Infanticide more likely during low prey availability | - |
|  | Infanticidal individuals consume infant | +/- |
| Resource competition:<br>Breeding space | Infanticide more likely when many cubs at communal den | - |
| Resource competition:<br>Social status | Infanticide committed by high-ranking individuals | + |
|  | Infanticide perpetrators and victims more likely philopatric sex (female) | +/- |

Note. Four hypotheses about the evolution of infanticide in hyenas, the predictions made by these hypotheses, and whether our observations support (+) or fail to support (-) each prediction.

Here we use three decades of behavioral observations from multiple social groups to provide the first quantitative assessment of the prevalence and context of infanticide in spotted hyenas, a plural breeding species with female philopatry and nepotistic rank inheritance. We first examine the biological significance of infanticide in this species by quantifying its contribution to overall mortality. We then describe the contexts in which infanticide occurs. Finally, we evaluate the predictions of these four hypotheses regarding the evolution of infanticide in spotted hyenas.

## Methods

### Study Animals

Spotted hyenas are large carnivores found widely across sub-Saharan Africa (Holekamp & Dloniak, 2010). Individuals reside in mixed-sex clans, each of which contains multiple matrilineal kin groups and is structured by a linear dominance hierarchy (Frank, 1986). The dominance hierarchy is maintained by social support from groupmates, especially kin (Smith et al., 2010; Strauss & Holekamp, 2019b; Vullioud et al., 2019), and rank is inherited through a learning process akin to what is found in many cercopithecine primates (Holekamp & Smale, 1991). Spotted hyena societies are characterized by fission-fusion dynamics where individuals associate in subgroups that change composition throughout the day (Smith et al., 2008). Males disperse during the years after reproductive maturity, which occurs at around 24 months of age (Holekamp et al., 2012). Spotted hyenas are polygynandrous and breed year-round. Females give birth to one or two (and rarely, three) cubs in an isolated natal den, where they are maintained for a few weeks before being moved to the clan’s communal den. The communal den may contain up to 31 cubs at any given time (Johnson-Ulrich & Holekamp, 2020), and cubs typically remain in or around the den until they are 9-12 months of age (Holekamp & Dloniak, 2010). These cubs, which belong to several different mothers, are often left unattended during much of the day while the mothers are away. Both mothers and other groupmates visit the communal den regularly, either alone or with clan-mates. Starting at 1-2 months of age, cubs emerge from the den to socialize when their mothers are present and, as they get older, when their mothers are absent.

### Study Area

Data presented here were collected from two study areas in Kenya near the Tanzanian border. Most observations come from eight clans in the Masai Mara National Reserve (MMNR, 1510 km^2^), a savanna ecosystem in southwestern Kenya that is contiguous with the Serengeti National Park in Tanzania and grazed year-round by multiple herbivore species (Holekamp et al., 1997). Data were collected in MMNR from 1988 to 2018 during 149,377 observation sessions; we observed one clan for the entire study period, and seven other clans during subsets of that period. Our second study area was in Amboseli National Park (ANP, 392 km^2^), which is located in southeastern Kenya; data were collected from two clans in ANP from 2003 to 2005 during 4,651 observation sessions (Watts et al., 2011).

For describing cases of infanticide and their contribution to mortality, we use data from all ten clans of spotted hyenas located in both study areas to capture the full breadth of circumstances under which infanticide occurs. For tests of hypotheses about the evolution of infanticide, we limited the dataset to only our four most well-studied clans of hyenas in the MMNR. We elected to use this restricted dataset because these clans live under similar ecological conditions, because all covariate data were available for these clans, and because these clans account for the majority of our data (73% of juvenile mortality from the larger dataset).

### Data Collection and Analysis

Data were collected during twice-daily observation periods that took place around dawn and dusk. Observers used vehicles as mobile blinds from which to find and observe hyenas. Scan sampling (Altmann, 1974) was used to collect demographic data. Maternity was determined based on nursing associations and genotyping. Data documenting specific types of social interactions, including observed infanticide events, were collected using all-occurrence sampling (Altmann, 1974). Individual hyenas were identified by their unique spot patterns, and the sexes of juveniles were determined based on the dimorphic morphology of the erect phallus (possible after cubs are 2-3 months old). Juvenile age was determined (to ± 1 week) based on appearance when first seen. All statistical analysis and visualization was done using the statistical software environment, R (R Core Team, 2020).

### Juvenile mortality

We assessed various causes of mortality among juveniles less than 1 year of age. To do this, we monitored all births in our study groups (n = 1643) and identified mortality events when a juvenile disappeared or when we directly observed mortality. In our 30 years of data, we documented 705 cases of juvenile mortality (43% of total births), and in 102 cases (15% of mortality) we were able to determine a cause of mortality. We observed five typical mortality sources for young juvenile hyenas: starvation, humans, lions, siblicide, and infanticide. Starvation was identified in cases where juveniles were observed to be becoming progressively gaunter before they vanished, or in cases where dead juveniles were found in an emaciated state. Death by humans was assigned when there was a clear anthropogenic cause of mortality; for example, hit by a car, speared, poisoned, etc. Death by lions was either observed or could be inferred based on the deep, widely spaced puncture wounds typically inflicted during lion attacks. Death by siblicide occurred when a cub prevented its littermate from nursing (Golla et al., 1999; Smale et al., 1999).

We identified 21 cases of infanticide in our dataset, divided into two categories based on our confidence in the cause of death. We identified 17 ‘confident’ cases of infanticide, where we observed a cub being killed by another hyena by having its skull crushed or found a cub dead as a result of having its skull recently crushed (Figure 1). Additionally, we identified four ‘likely’ cases of infanticide, where a dead cub was found but no information on the state of the skull was available (usually because the victim had already been largely consumed), but other common aspects of ‘confident’ cases of infanticide were noted (e.g., mother guarding or grooming the victim, victim consumed by mother or others, victim found at communal den). Finally, cubs dying due to causes other than those listed here were combined into an ‘other’ mortality source.

**Figure 1.**
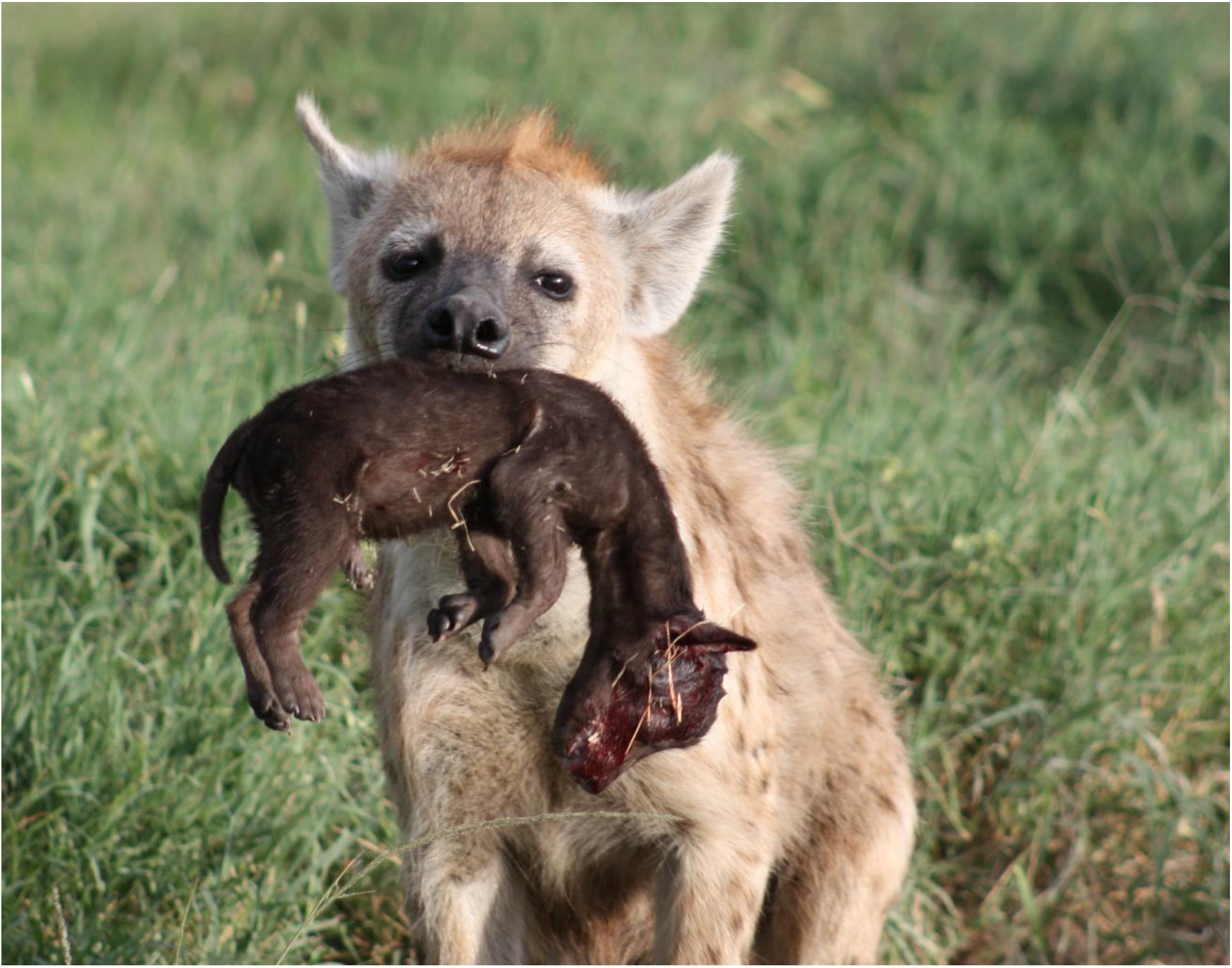
Adult female spotted hyena carrying a cub recently killed by infanticide. Infanticide was typically achieved by crushing the skull, as seen here. Photo by Kate Yoshida.

### Determining contribution of infanticide to overall mortality

We next examined the extent to which infanticide contributes to juvenile mortality compared to other mortality causes. The first step in this process was to account for the occurrence of orphaning. We separated mortality into cases where the cub was orphaned before death and cases where the cub perished with a living mother. We did this because cubs under 12 months old whose mothers die also perish in almost all cases. These juveniles are still dependent on their mother’s milk and therefore 81% of juvenile deaths in this category are a result of starvation. In the remaining 19%, although cause of death was not directly starvation (causes of death were humans [n=3], infanticide [n=1], and lions [n=3]), prolonged absence of the mother combined with nutritional stress likely motivated the cubs to engage in abnormal behavior. To account for these cases, we re-classified these known mortality sources to be ‘death of mother.’ For orphaned cubs with unknown cause of death (n = 38), we inferred their mortality source to be ‘death of mother.’ Together, this known and inferred mortality due to orphaning made up 11% of juvenile mortality.

We next explored the contributions of different mortality sources to the remaining 89% of mortality cases, where juveniles perished with living mothers. Because most juvenile mortality in the dataset had unknown causes, and because mortality of different types was non-random with respect to age (see Results), determining the contributions of different sources of mortality was not as simple as adding up all observed cases. Instead, we used differences in age-distributions of known mortality sources to partition mortality of unknown cause into likely sources. We did this using a two-step modeling process to estimate the overall contribution of different mortality sources to this subset of juvenile mortality: we (1) modeled known mortality source as a function of age at death and (2) used this model to partition mortality with unknown cause into likely causes. To do this, we built a Bayesian multinomial model of mortality source for the remaining juveniles with living mothers (n = 66) as a function of their age at death. We assessed the efficacy of this model by using leave-one-out cross validation to compare it to a null model without age at death as a predictor (Vehtari et al., 2017). We then used 200 posterior samples from this model to generate 200 predictions for the cause of death for each juvenile with an unknown mortality source (n = 565). In other words, for each juvenile with unknown mortality, we used the model to generate 200 predicted mortality sources based on the age of death of the juvenile, and these predictions were used to estimate the mean (with 95% prediction intervals) contributions of each mortality source to unknown mortality. The overall contribution of each mortality source was thus estimated as the sum of these posterior means from the multinomial model and the number of observed cases with known mortality due to each cause.

### Evaluating hypotheses about the evolution of infanticide

To test whether infanticide was more likely to occur when prey were scarce, we compared the average prey density in the study group of each individual using prey transect surveys. Twice each month we drove a designated prey transect and counted all prey animals within 100m of the vehicle, then divided by the total area covered by the transect to calculate the density of prey animals in the clan. For each cub with a known mortality source, we calculated the average prey density in all transects conducted in the cub’s study group in the month prior to death. We then used a multinomial model to model mortality source as a function of prey density (Supplemental Materials). We evaluate the model by comparing it to a null model without mortality source using leave-one-out cross validation (Vehtari et al., 2017).

To test whether infanticide was more likely to occur when there were many cubs using the communal den, we measured the density of den-dwelling cubs in the month prior to death. For each cub with a known mortality source, we calculated the average number of cubs with which they were observed over all observation sessions in the month prior to death. We again used a multinomial model to model mortality source as a function of cub density (Supplemental Materials). We evaluate the model by comparing it to a null model without mortality source using leave-one-out cross validation (Vehtari et al., 2017).

All three multinomial models were implemented in Stan using the *rstan* and *brms* R packages (Bürkner, 2017; Stan Development Team, 2018). We assessed model convergence using the potential scale-reduction factor (R-hat), which was required to be below 1.1 (Gelman & Rubin, 1992). See Supplemental Materials for model details and diagnostics.

To test whether killers and mothers of victims differed in rank, we first determined ranks of adult females based on the outcomes of aggressive interactions as described elsewhere (Strauss, 2019; Strauss & Holekamp, 2019a). Rank is standardized to account for group size so that it ranges from 1 (highest rank) to −1 (lowest rank). We then compared the ranks of killers with the ranks of victims using an unpaired Welch’s two-sample t test.

## Results

### Quantifying the contribution of infanticide to overall mortality

Infanticide was a leading source of mortality for cubs under one year old. Of the 102 cub deaths with known mortality source (of 705 total juvenile mortality cases), infanticide accounted for 20% of mortality (Figure 2, dark bars). This number places infanticide as the second largest mortality source for den-dependent juveniles, ranking below death of mother (35%) but above lions (15%) (Figure 2, dark bars). Mortality sources were age-structured, with the model including age at death as a predictor outperforming the null model without the predictor in leave-one-out cross validation (elpd = 16.2, SE = 6.2). The multinomial model revealed that cubs who died young were most likely victims of infanticide, whereas cubs who died at ages over 4.2 months were most likely victims of lions (Figure 3). After using this model to predict cause of death for cubs dying due to unknown cause (Figure 2, light bars), infanticide and lions were the leading causes of death for juveniles, respectively accounting for a mean of 24% (95% prediction interval = [18% - 30%]) and 24% [18%, 31%] of juvenile mortality. Death of mother, starvation, siblicide, humans, and other mortality sources together accounted for 52% of total juvenile mortality (Figure 2). This analysis estimates that 10% (95% PI: 8% - 13%) of all hyenas born in our population are killed by infanticide.

**Figure 2.**
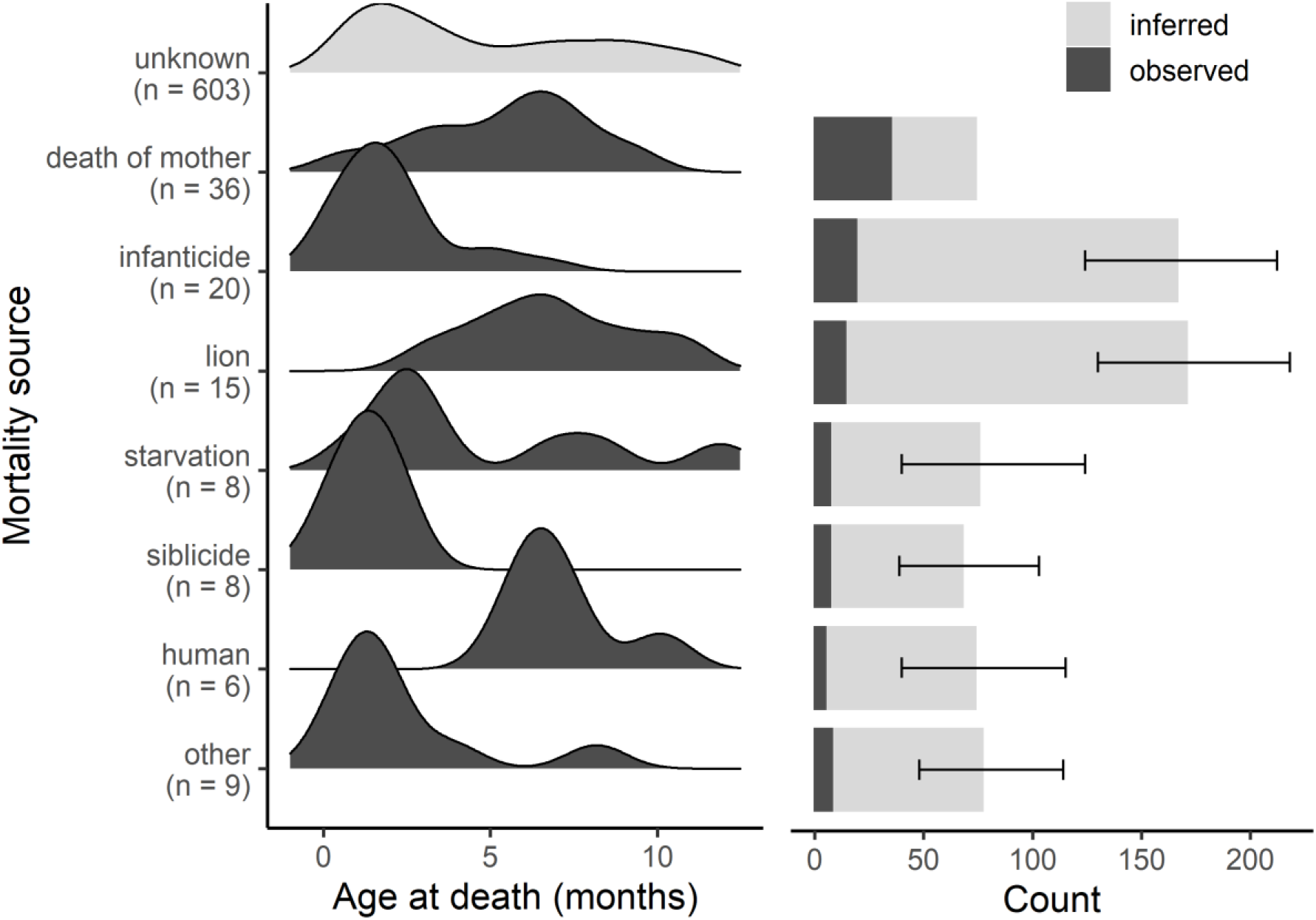
Age distribution of observed causes of juvenile (<1-year-old) mortality in spotted hyenas (left), and frequencies of the 6 leading causes of juvenile mortality (right). Dark bars indicate cases where the mortality source was known (n = 102). Lighter bars indicate inferred mortality sources for cases where the cause of mortality was unknown (n = 603). Error bars indicate the 95% prediction intervals for inferred mortality cases based on a Bayesian multinomial model of mortality source as a function of age. Mortality by ‘death of mother’ was inferred analytically rather than statistically.

**Figure 3.**
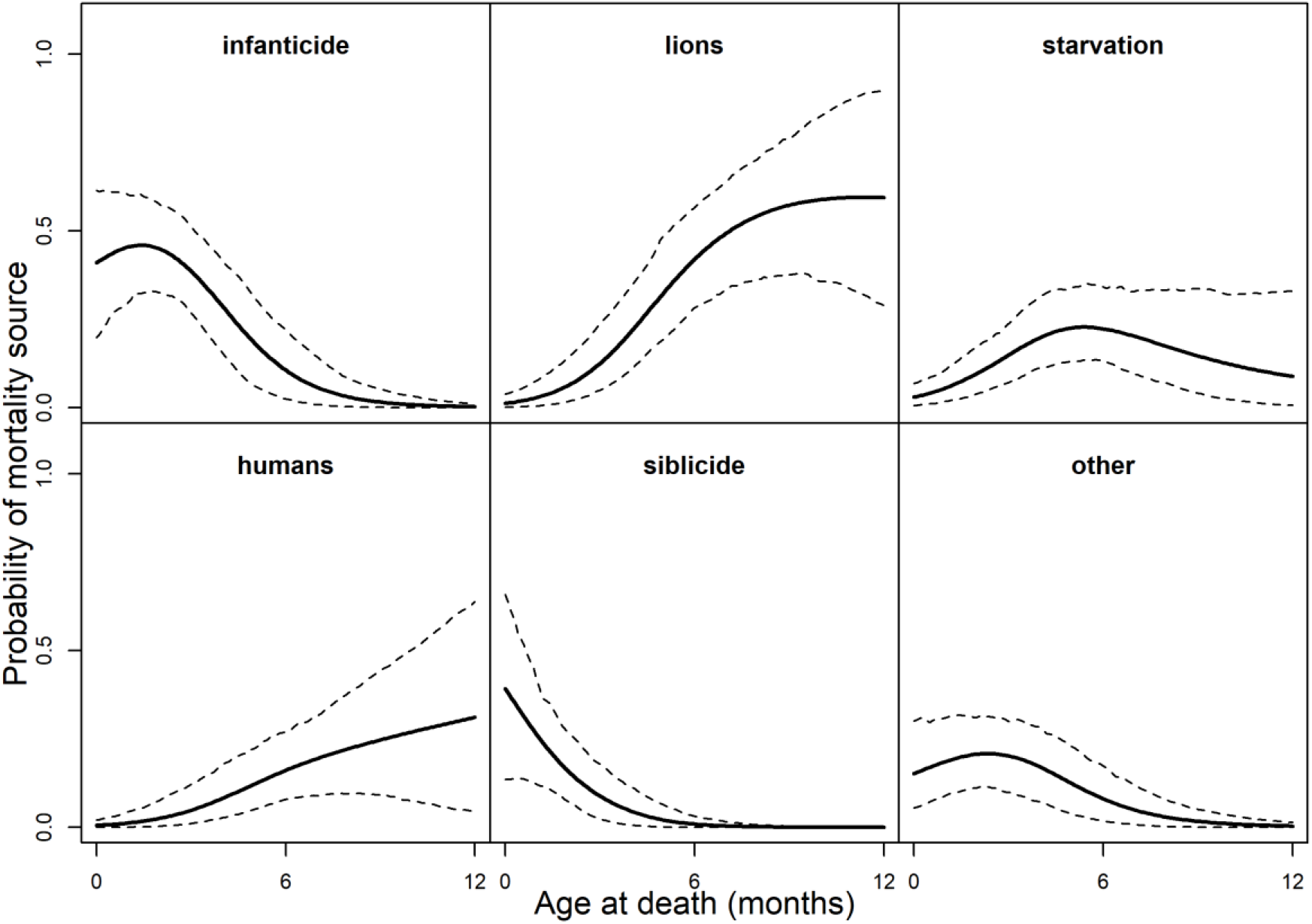
Probability of different mortality sources as a function of age at death, as estimated by a Bayesian multinomial model. Juveniles under 4.2 months old are most at risk of death by infanticide, whereas older juveniles are most at risk of death by lions.

### Describing the contexts in which infanticide occurs

In every case of infanticide with a known killer, the act was perpetrated by an adult female (n = 10). Infanticidal females typically targeted young juveniles (<5 months old), although we did observe two cases of infanticide among older juveniles (Figure 2). Victims with known sexes were evenly split between males (n = 4) and females (n = 5), suggesting that juveniles of both sexes were equally likely to be attacked by conspecifics.

All cases of infanticide occurred at a communal den, although in one case the victim was killed by other groupmates as its mother was transferring it from the natal den to the communal den. Attackers sometimes acted alone (n = 7), and other times were aided by groupmates (n = 3). In three cases where infanticide took place while the victim’s mother was present, multiple hyenas displayed aggression against the mother while her offspring was being attacked. In one of these events, the highest-ranking female killed a low-ranking juvenile while her offspring chased away the victim’s mother. In two other cases where the victim’s mother was present, multiple hyenas attacked the mother while the perpetrator killed the cub. In cases where females committed infanticide unaided, they often did so during what appeared to be normal social behavior, and in a few cases prosocial ‘groan’ vocalizations were emitted by the attacker immediately before attacking. Close kinship did not prevent females from committing infanticide: in one case, a female coaxed each of her full sister’s two offspring out of the den by groaning, then killed both cubs (previously reported in (White, 2005)). However, although we did not have full pedigree data available for most killed infants, infanticidal females most often killed juveniles other than those born to their closest relatives. Thus, this prior report of infanticide against kin was not representative of typical patterns of infanticide in this species.

Victims’ bodies were consumed by one or more hyenas in 11 out of 21 cases; they were sometimes consumed by the killer (n = 3) or the killer’s offspring (n = 3), sometimes by the mother of the dead infant (n = 4), and sometimes by other group-members (n=3; note that these numbers do not add to 11 because multiple hyenas were often observed consuming infanticide victims). Consumption of deceased conspecifics was not restricted to infanticide - the remains of juveniles dying of other causes were also sometimes consumed, and hyenas are known to consume deceased adult conspecifics as well (Kruuk, 1972). When given the opportunity, mothers sometimes (n = 7) groomed, guarded, or otherwise cared for their infanticide victim for up to two hours after its death (Supplemental Material). Posthumous care and consumption by the mother were not mutually exclusive events, as we observed the mother eat her cub after grooming the dead body in 2 cases. In 3 cases, observers noted unusual, unprovoked, and distressed-sounding vocalizations emitted by the mother. In 5 cases, the body was either carried away from the den or completely consumed in less than 50 minutes, with one cub being completely consumed in 13 minutes. These records may underestimate the frequency with which victims are consumed because observers collected the victim’s body for biological samples in 6 cases, and halted observations before the fate of the body was determined in 4 cases.

### Evaluating hypotheses about the evolution of infanticide

We found differential support for the four hypotheses regarding the evolution of infanticide in spotted hyenas. These results come from a combination of observations of infanticide events and targeted tests of the predictions made by these hypotheses (Table 1). Neither prey density (n = 85) nor the number of cubs at the den (n = 80) varied significantly based on mortality source (Figure 4) – in both cases, leave-one-out cross validation suggested that null models were as good as or better than the models including prey density (elpd = 5.5, SE = 3) or cub density (elpd = 2.1, SE = 3.2) as a predictor. Furthermore, the parameter estimate for infanticide overlapped the 95% credible intervals surrounding parameter estimates for all but one of the other mortality sources in both the “prey density” and “number of cubs” models (Supplemental Material). We found significant differences between the average rank of killers and the average rank of the mothers of victims -- adult females were on average higher ranking than the mothers of victims (Welch’s two-sample t-test: t = −3.54, df = 21.89, p = 0.002; Figure 5). Whereas victims of infanticide were of diverse ranks, perpetrators of infanticide were almost exclusively high-ranking females.

**Figure 4.**
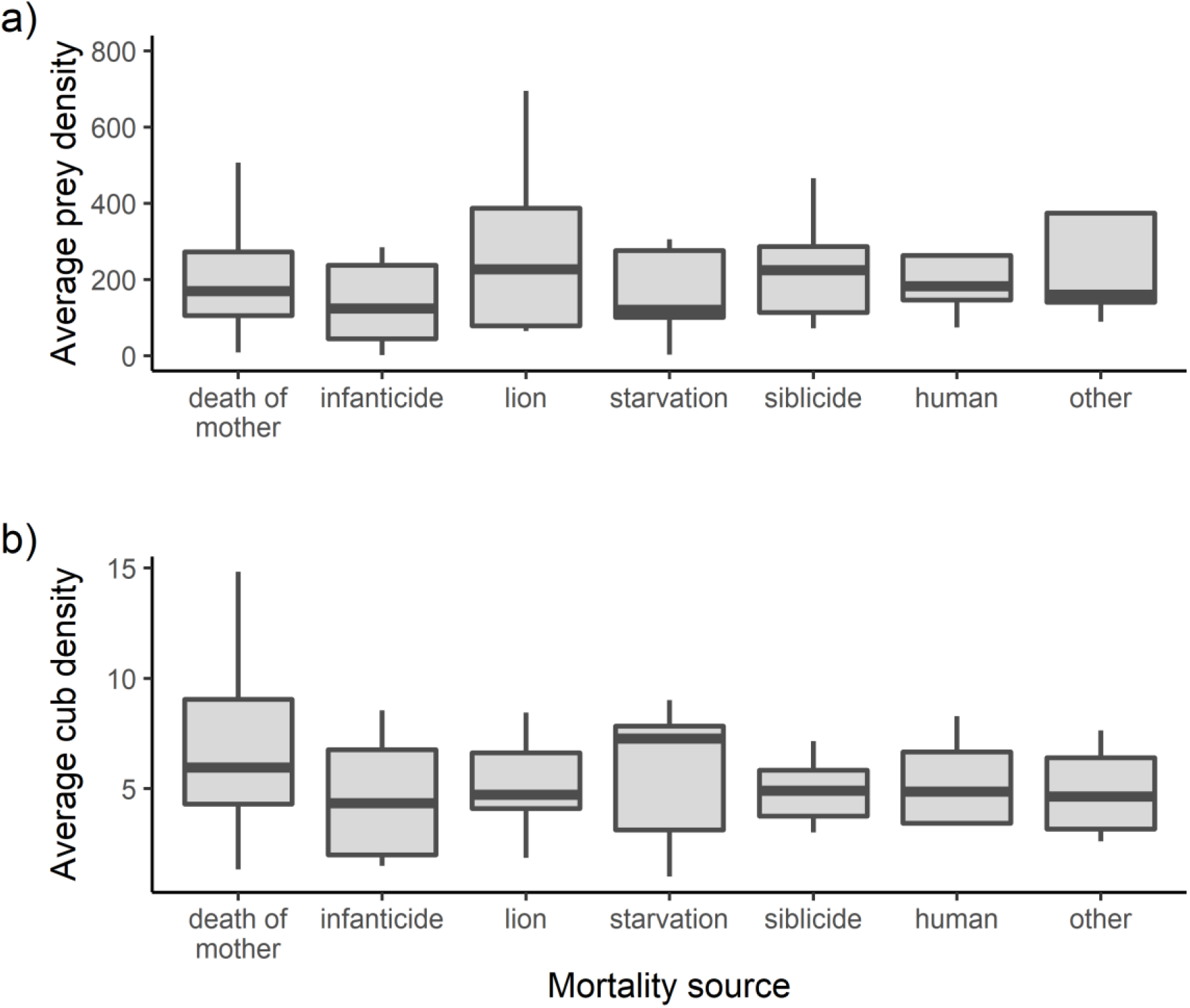
Differences in average a) prey density and b) cub density in the month prior to death by the leading causes of mortality. Neither prey density nor the number of cubs alive in the social group varied significantly by mortality source.

**Figure 5.**
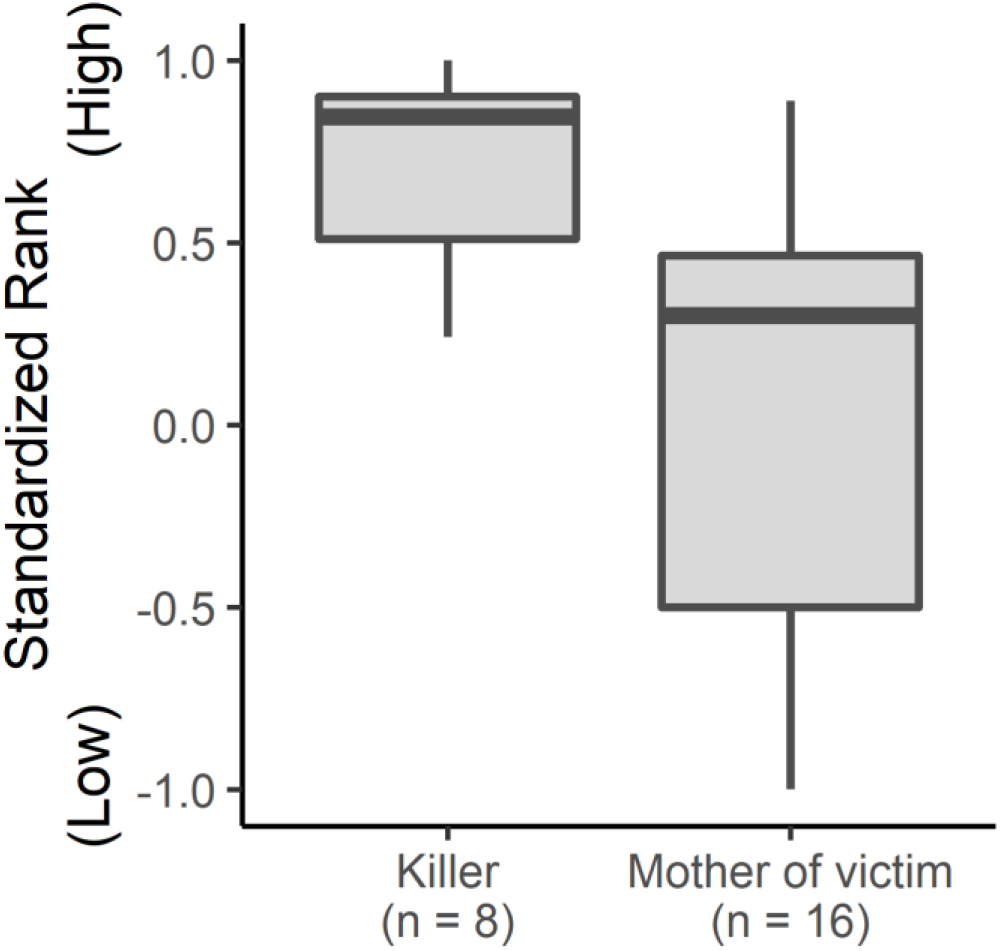
Rank distributions of killers and mothers of cubs killed by infanticide. Ranks of infant-killers were on average higher than the ranks of mothers of infants killed by infanticide (Welch’s two-sample t-test: t = −3.54, df = 21.89, p = 0.002).

## Discussion

Our results indicate that infanticide is a significant source of mortality among young juvenile spotted hyenas (Figure 2). Overall, there was significant age structuring in mortality sources for juveniles over their first year of life, where juveniles dying young were most likely victims of infanticide and juveniles dying at older ages were most likely killed by lions. Our observations most strongly support the competition over social status hypothesis for the evolution of infanticide in spotted hyenas. In support of predictions about infanticide in nepotistic societies (Clutton-Brock & Huchard, 2013; Lukas & Huchard, 2019; Vullioud et al., 2019), we found that infanticide was typically perpetrated by high-ranking individuals against the offspring of lower-ranked groupmates. The prediction of the social status hypothesis about the sex of perpetrators and victims was only partially supported – females were the only sex to commit infanticide but were not a more frequent target of infanticide. Contrary to the predictions of the sexual selection hypothesis, no males were documented committing infanticide in this study. The exploitation hypothesis received only middling support. Killers did sometimes consume the victims of infanticide, but they often chose instead to leave the body to be consumed by other clan members, and infanticide was not more associated with low prey density than other mortality sources. The prediction made by the breeding space hypothesis – that infanticide will be more associated with high densities of cubs - was also not confirmed.

Social support has been found to be a significant force for establishing, maintaining, and changing dominance in spotted hyenas (Engh et al., 2000; Strauss & Holekamp, 2019b; Vullioud et al., 2019), and kin provide a significant portion of that support (Smith et al., 2010). Thus, reducing another female’s matriline size via infanticide may serve to prevent potential coalitions of lower-ranked matrilines aspiring to improve their status. Interestingly, we did observe one case of infanticide directly related to escalated aggression between matrilines. Observers arrived at the den to find a recently killed offspring from a high-ranked matriline. Many hyenas were acting highly agitated, and roughly one hour later we observed a coalition of related low-ranking adult females viciously attacking members of the high-ranking matriline. Our observations of infanticide events where the perpetrator’s kin assisted by chasing away the victim’s kin reflects how infanticide might both arise as a function of disparities in social support within groups and serve to reinforce those disparities.

Interestingly, our findings suggest that many of the infanticide anecdotes that have previously been published are not representative of infanticide in this species. Contrary to what was suggested by Kruuk (1972) and East et al. (2003), infanticide in our study was never perpetrated by males. We also found no support for the suggestion that infanticide is committed by members of other social groups, as suggested by Mills (1990). We were not able to test Muller & Wrangham’s (2002) hypothesis about a potential infanticide-avoidance function to the peniform clitoris of spotted hyenas. We found that males and females were equally likely to be targeted by infanticide, which could indicate either that females faced the same risk of infanticide as males (contrary to Muller and Wrangham’s hypothesis), or that the peniform clitoris was successful at reducing this risk (in support of Muller and Wrangham’s hypothesis).

Our findings also indicate a dual function of the communal den as protection against both outside predation sources such as lions, and intraspecific killing via infanticide. The diameter of den holes limits the size of individuals able to enter the den, such that cubs can escape inside when threatened by adults or large predators too large to fit into the den hole. This function has been born out in our observations: we have seen larger cubs killed by lions while they attempted escape into the communal den, as well as cubs escaping into the den while lions attempt (and fail) to extract them with their paws (unpublished data). In our observations of infanticide, we occasionally observed perpetrators coaxing cubs out of the den before attacking them, and in one case a targeted juvenile attempted to escape into the den but was caught and killed before reaching safety (Supplemental Videos). Mothers of victims sometimes displayed grooming or other maternal behaviors towards the deceased cub, which may be of interest to those interested in comparative thanatology (Anderson & Anderson, 2016; Carter et al., 2020).

Our results highlight the conflicting forces that characterize the lives of gregarious animals. The prevalence of infanticide highlights risks faced by females choosing to rear their cubs in a social environment. More solitary individuals could choose to keep their cubs at a natal den for several months, and thereby avoid the potential risks of infanticide, but female hyenas rarely choose to do so (White, 2007). This suggests that the benefits of social integration for cubs raised at communal dens outweigh the costs to females imposed by the risk of conspecific infanticide. However, these conflicting forces may also lead to a tradeoff between appropriate social development and survival if the behavior required for social integration in early life is associated with infanticide risk.

Overall, our results suggest strong selection in this species for behaviors that mitigate infanticide risk and thus alleviate the tradeoffs of social reproduction. For instance, females may mitigate infanticide risk by timing den visits to avoid infanticidal groupmates or to coincide with the presence of kin who may help protect against infanticide. Future work should aim to identify potential behaviors that reduce the risk of infanticide and their links to survival.

Finally, our results demonstrate the power of using long-term data to study rare or difficult-to-observe phenomena. Rarity of phenomena can obscure their importance, and ephemeral, high-impact events like infanticide are the types of phenomena that require extensive data collection to permit analysis. The frequency of infanticide relative to other sources of infant mortality suggest that it is a significant feature of spotted hyena biology, which was unclear prior to this study. Continued, direct study of this phenomenon has the potential to answer outstanding questions about the function of infanticide and conflicts of interest within complex societies.

## Acknowledgments

We thank the Kenya Wildlife Service, the Narok County Government, and the Kenyan National Committee on Science, Technology and Innovation, the Naboisho Conservancy, the Mara Conservancy and Brian Heath for permissions to conduct this research. Thanks to Maggie Sawdy for detective work about the identities of some juveniles, and many current and former members of the Mara Hyena Project for detailed data collection. We also thank two anonymous reviewers for their insightful suggestions on the manuscript. This work was supported by NSF grants OISE1853934 and IOS1755089 to KEH, and by an NSF Graduate Research Fellowship to EDS. This work was also supported in part by NSF Grant OIA 0939454 (Science and Technology Centers) via “BEACON: An NSF Center for the Study of Evolution in Action.” AKB was supported by the MSU Honors College.

## Statement of Authorship

The data were collected and validated by MOP, KEH, and EDS. Data processing, analysis, and visualization were done by AKB and EDS. AKB, KEH and EDS wrote the original draft of the manuscript. AKB, MOP, KEH and EDS edited the manuscript.

## Data and Code Accessibility

Data and code used for this study are available on GitHub (https://doi.org/10.5281/zenodo.4889918) and on Dryad Digital Repository: https://doi.org/10.5061/dryad.0zpc866xm (Brown et al., 2021).

